# Translocation of dense granule effectors across the parasitophorous vacuole membrane in *Toxoplasma*-infected cells requires the activity of ROP17, a rhoptry protein kinase

**DOI:** 10.1101/613208

**Authors:** Michael W. Panas, Abel Ferrel, Adit Naor, Elizabeth Tenborg, Hernan A. Lorenzi, John C. Boothroyd

## Abstract

*Toxoplasma gondii* tachyzoites co-opt host cell functions through introduction of a large set of rhoptry- and dense granule-derived effector proteins. These effectors reach the host cytosol through different means: direct injection for rhoptry effectors and translocation across the parasitophorous vacuolar membrane (PVM) for dense granule (GRA) effectors. The machinery that translocates these GRA effectors has recently been partially elucidated, revealing 3 components, MYR1, MYR2 and MYR3. To determine if other proteins might be involved, we returned to a library of mutants defective in GRA translocation and selected one with a partial defect, suggesting it might be in a gene encoding a new component of the machinery. Surprisingly, whole-genome sequencing revealed a missense mutation in a gene encoding a known rhoptry protein, a serine/threonine protein kinase known as ROP17. ROP17 resides on the host-cytosol side of the PVM in infected cells and has previously been known for its activity in phosphorylating and, thereby, inactivating host immunity-related GTPases. Here, we show that null or catalytically dead mutants of ROP17 are defective in GRA translocation across the PVM, but that translocation can be rescued “in *trans"* by ROP17 delivered by other tachyzoites infecting the same host cell. This strongly argues that ROP17’s role in regulating GRA translocation is carried out on the host-cytosolic side of the PVM, not within the parasites or lumen of the parasitophorous vacuole. This represents an entirely new way in which the different secretory compartments of *Toxoplasma* tachyzoites collaborate to modulate the host-parasite interaction.

**Importance:** When *Toxoplasma* infects a cell it establishes a protective parasitophorous vacuole surrounding it. While this vacuole provides protection, it also serves as a barrier to the export of parasite effector proteins that impact and take control of the host cell. Our discovery here that the parasite rhoptry protein, ROP17, is necessary for export of these effector proteins provides a distinct, novel function for ROP17 apart from its known role in protecting the vacuole. This will enable future research into ways in which we can prevent the export of effector proteins thereby preventing *Toxoplasma* from productively infecting its animal and human hosts.

## Introduction

*Toxoplasma gondii* is an obligate intracellular parasite capable of infecting a wide range of cell types in almost any warm-blooded animal. As for most *Apicomplexa,* entry into a host cell and interaction with host functions once inside involves the coordinated action of at least three distinct secretory compartments: micronemes, rhoptries and dense granules [1]. The small, apical micronemes are the first to function in invasion, releasing adhesins onto the surface of the parasite that are crucial for attachment to the host cell [2,3]. The much larger, bulb-shaped rhoptries are also apically located and these somehow directly introduce their contents into the host cell at the start of actual invasion [4]. Once initiated, invasion involves an invagination of the host plasma membrane to form a parasitophorous vacuole (PV). This process is mediated by binding between a surface-localized micronemal protein, AMA1, and RON2, a protein that starts in the rhoptry necks (hence “RON”) but is introduced into the host cell to become an integral membrane protein within the host plasma membrane [5–7].

In addition to the RON proteins, rhoptries also introduce the contents of their bulbs during invasion [8,9]. These proteins, generally known as ROPs, are a set of effectors whose job generally appears to be to co-opt host functions [4,10–12]. Most known ROPs are members of an extended family of protein kinases and pseudokinases defined by their homology to the prototypical member of the family, ROP2 [13]. Among the active ROP2-like kinases are ROP17 and ROP18 which have been well studied for their role in defense against immunity-related GTPases, a set of host proteins that are generated in response to interferon-gamma and that attack the parasitophorous vacuole membrane (PVM) resulting in eventual death of the parasites inside [14–17]. In collaboration with the pseudokinase ROP5, ROP17 and ROP18 phosphorylate IRGs which disrupts their ability to bind GTP, thereby neutralizing their ability to attack the PVM [18–21]. The location of these ROPs at the PVM [22,23], specifically on the host-cytosolic side of this membrane [24], perfectly positions them for their role in defending against IRG attack. In addition, ROP17 has been shown to impact the host transcriptional network directly by as yet unknown means [25] and to be essential for full virulence in mice [26].

A third secretory compartment, the dense granules, also plays a key function in the interaction with the host cell. The contents of these spherical organelles are known as GRAs and they are released into the PV after invasion is underway [27]. Unlike rhoptry proteins, however, GRAs are not injected directly into the host cytosol but instead are secreted into the PV space [28–30]. Some GRA proteins are involved in elaboration of the PVM into a complex network of nanotubes known as the intravacuolar network [31,32]. Others associate with or even integrate into the PVM where they mediate a variety of host functions including recruitment of host mitochondria [33] and activation of host NFκB [34]. A third set of GRA proteins including GRA16, GRA18, GRA24, and TgIST, however, are translocated across the PVM and into the host cytosol, with some eventually reaching the host nucleus where they have a profound effect on many host functions [11,35]. This class of GRA proteins impact the activity of host p53 [36] p38 MAPKinase [37], STAT1 signaling [38,39], beta-catenin signaling [40], E2F signaling [41], and c-Myc expression [42].

Using a genetic screen for *Toxoplasma* genes necessary for the aforementioned host c-<u>My</u>c upregulation, we have previously demonstrated that the translocation of GRAs across the PVM involves a set of parasite proteins that originate in dense granules or dense-granule-like organelles, ultimately reaching the PVM [42,43]. These MYR (<u>My</u>c Regulation) genes were identified by using fluorescence-activated cell sorting (FACS) to select from a population of chemically mutagenized *Toxoplasma* tachyzoites those mutants that fail to upregulate a GFP-c-Myc reporter fusion in bone marrow macrophages. Whole-genome sequencing of clones from the resulting populations of *Toxoplasma* mutants revealed three novel genes as necessary for the c-Myc upregulation, *MYR1, MYR2* and *MYR3* [42,43]. MYR1 and MYR3 form a stable complex at the PVM [43] and it is presumed that these two proteins are part of a translocon system that mediates the movement of GRAs across this membrane. MYR2 is also at the periphery of the PV but it has not yet been found to associate with either of the other two MYR proteins. Deletion of any one of the three *MYR* genes results in a complete loss of GRA translocation and, as expected for a mutant that cannot introduce an entire class of crucial effector proteins, Δ*myr1* strains have a much-reduced impact on the host transcriptome [41]; they are also substantially attenuated in a mouse model of virulence [42].

*A priori,* it seemed likely that more than just these three proteins would be necessary for the translocation of GRA effectors across the PVM. To address this possibility, we returned to the original library of Myr^-^mutants [42] and asked if any of the mutants had a partial defect which might indicate that they were defective in a gene other than *MYR1/2/3.* We report here the isolation of one such mutant which was found to have a missense mutation in *ROP17* and go on to show that a functional ROP17 within the host cell is indeed necessary for the translocation of GRA proteins across the PVM, indicating that a rhoptry-derived kinase located at the PVM plays an unanticipated role in this crucial process.

## Results

We previously reported the use of a forward genetic screen to identify *Toxoplasma* genes necessary for the upregulation of mouse c-Myc expression [42]. This led to the identification of *MYR1, MYR2* and *MYR3,* mutants for all of which show a complete loss of GRA16-or GRA24-translocation across the PVM [42,43]. To determine if other genes might be involved, we returned to the mutant libraries and screened them for a different phenotype, *partial* loss of effector translocation. This was done by isolating 42 individual clones from the two libraries and assessing their ability to translocate HA-tagged GRA16 and MYC-tagged GRA24 to the host cell nucleus in infected human foreskin fibroblasts (HFFs). Most of the mutants obtained showed an essentially total loss of such translocation but one, clone MFM1.15, showed an intermediate phenotype (Fig. 1A, B).

**Figure 1.**
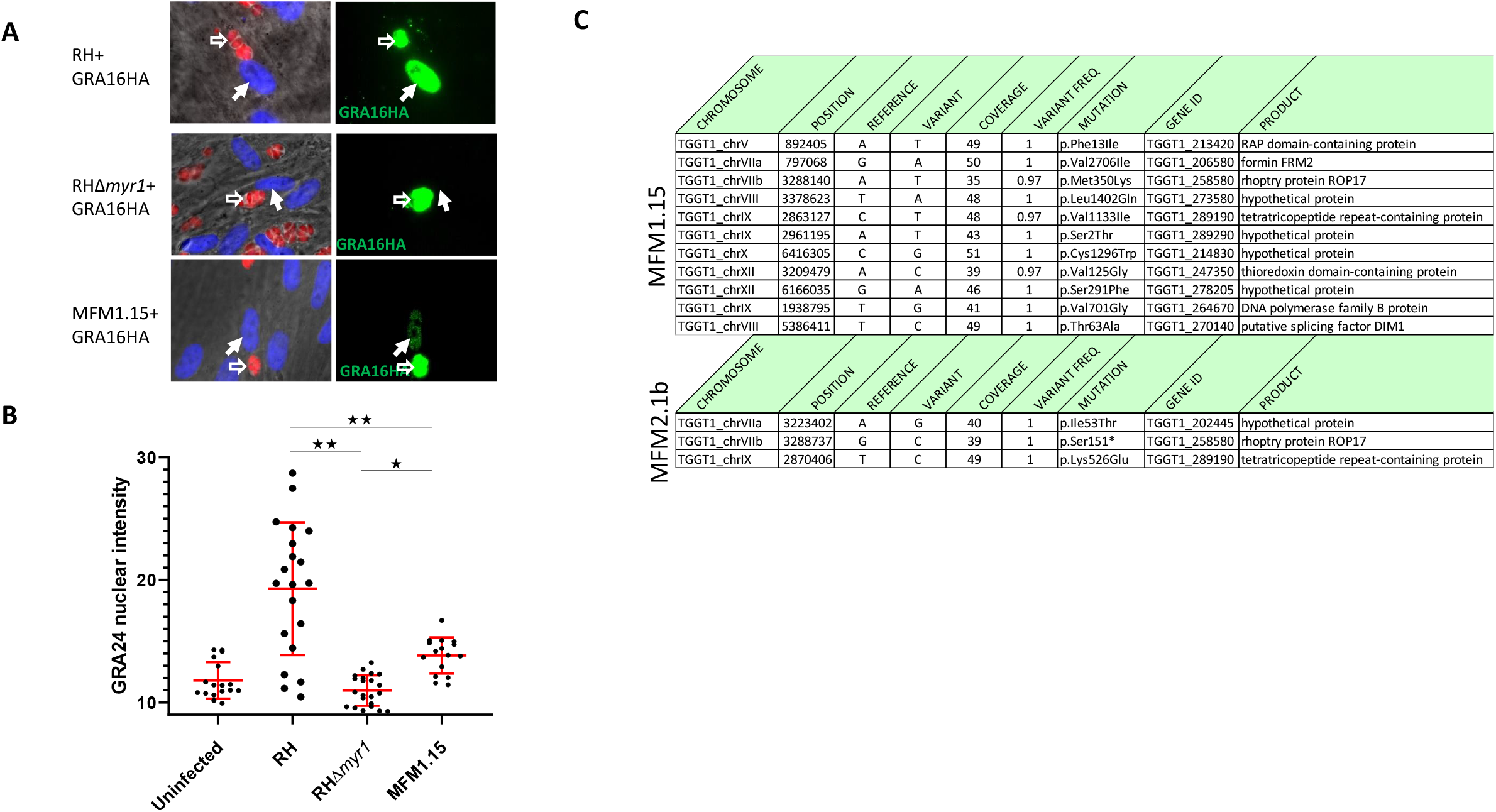
MFM1.15 shows a partial Myr^-^ phenotype and has a mutation in *ROP17*. **A**. Representative immunofluorescence assay (IFA) images of HFFs infected with RH-wild type, RHΔ*myr1* or RH mutant MFM1.15. All three parasite strains express cytosolic td-tomato (red) and were transfected with a plasmid expressing HA-tagged GRA16 which was detected by probing with anti-HA (green). Hollow arrows indicate vacuoles containing parasites expressing the GRA16HA transgene. Solid arrows indicate the nuclei in cells containing such vacuoles. Only the RH-WT-infected cells show efficient translocation of GRA16HA to the host cell nucleus. One of three biological replicates is shown. **B**. Translocation of GRA24 to the nucleus is also disrupted in mutant MFM1.15. Quantitation of nuclear GRA24MYC was assessed by anti-MYC tag staining followed by ImageJ to determine the intensity of nuclear staining in transiently transfected parasites from at least 10 random fields. The MFM1.15 mutant shows a significantly reduced nuclear signal but still more than the essentially complete lack of signal in the RHΔ*myr1* mutant. Error bars indicate standard error of the mean. *: p<0.05, **: p<0.0001. One of two biological replicates is shown. C. Mutations identified by whole genome sequencing of mutant MFM1.15 and the sub-clone MFM2.1b. Coverage is the number of reads spanning the indicated nucleotide and Variant Freq. is the fraction of reads showing the variant nucleotide relative to reference (GT1). Both mutants show mutations relative to the annotated Type I strain, GT1, in *TGGT1_258580 (ROP17)* with a missense mutation in MFM1.15 and a nonsense mutation in MFM2.1b.

The partial phenotype of MFM1.15 suggested that it might harbor a mutation partially inactivating a gene necessary for translocation, e.g. one of the *MYR* genes, or else it might harbor a mutation completely ablating expression of a novel gene that is only partly necessary for translocation and the c-Myc induction. To resolve this, we first used Sanger sequencing to confirm there was no mutation in the *MYR1, MYR2* or *MYR3* loci, and then subjected the clone to whole genome sequence analysis and identified the 11 mutations shown in Fig. 1C (upper panel). None of these mutations were in a known *MYR* gene but one stood out as being a missense mutation in a gene encoding a known protein kinase present at the PVM, ROP17. This raised the tantalizing possibility that ROP17 might play a role in the translocation of GRA proteins across the PVM. Consistent with this, we returned to previous datasets and saw that a nonsense mutation in *ROP17* had also been seen in one of the original screens that yielded MYR1 [42]. In this latter instance, the supposed “clone” that was sequenced, MFM2.1, turned out actually to be a pair of clones, such that all the mutations detected were present in only about 50% of the sequence reads. One of these mutations was in a gene that was mutated in two other (true) clones analyzed in the same set and so this gene was pursued and eventually shown to be essential for c-Myc upregulation and was thus designated *MYR1*. At the time, we did not know which of the other mutations detected were random “hitch-hiker” mutations in the *myr1* mutant vs. which might be key to the phenotype in the other clone present in the MFM2.1 pair. To resolve this, we recloned MFM2.1 parasites by limiting dilution and searched for mutants that had a wild type MYR1 gene. One of these was fully genome sequenced and the result was mutant MFM2.1.b which was found to have the ROP17 S151* mutation, consistent with a defect in ROP17 being the defect that produced the Myr^-^ phenotype (Fig. 1C, lower panel).

The finding that *ROP17* is mutated in both MFM1.15 and MFM2.1.b strongly suggested that a functional ROP17 protein might be necessary for the c-Myc upregulation. To test this, we generated a knock-out of *ROP17* in an otherwise wild-type *Toxoplasma* by disrupting the open reading frame of *ROP17 (TGGT1_258580)* with insertion of the *HXGPRT* gene (Fig. 2A), confirming this disruption by PCR of the locus (Fig. 2B), and assessing the translocation of known effectors in HFF cells infected with the resulting mutant. Preliminary results indicated that disruption of *ROP17* does indeed prevent the parasite from exporting GRA16HA and GRA24MYC from the parasitophorous vacuole into the host nucleus when these constructs were transiently expressed in RHΔ*rop17* tachyzoites (Fig 2C).

**Figure 2.**
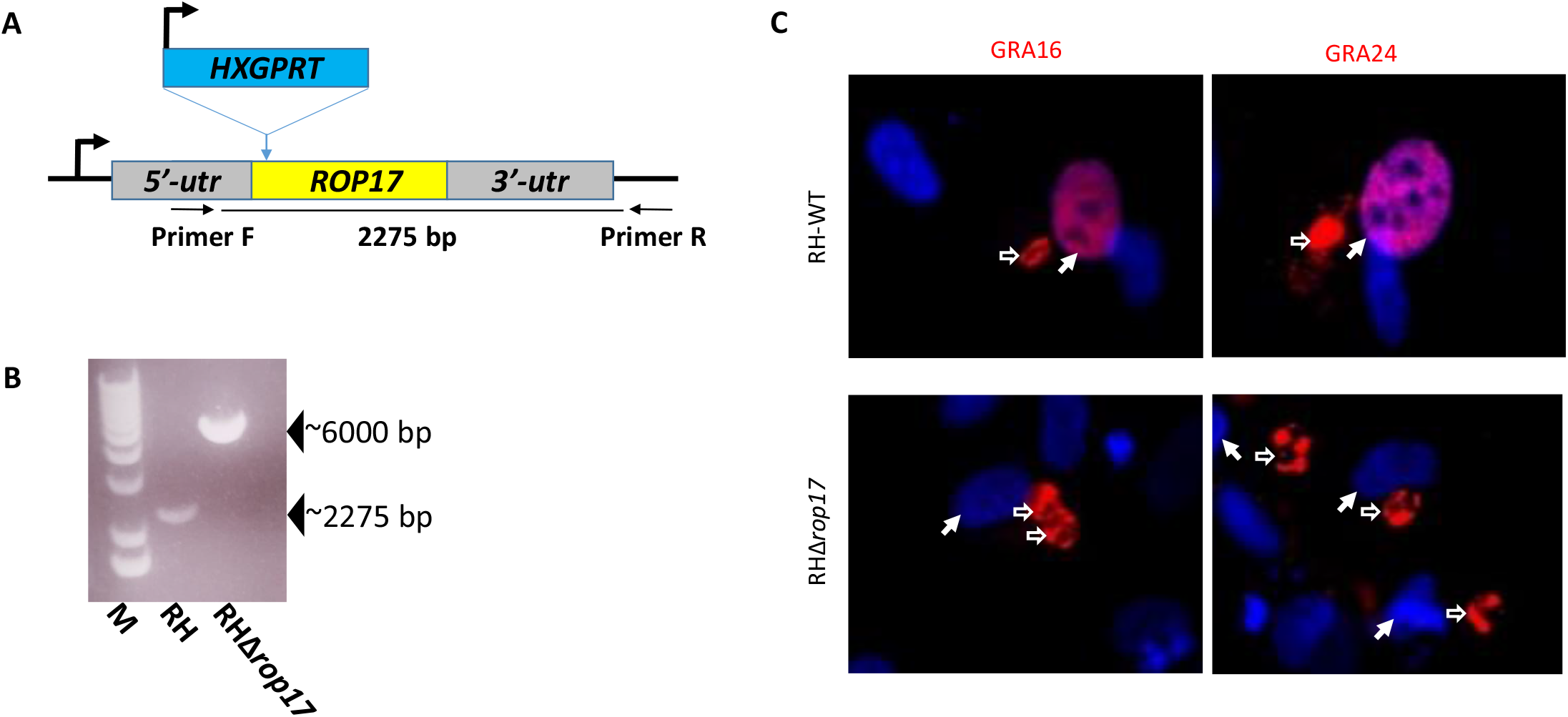
Deletion of *ROP17* generates a mutant parasite that cannot export GRA16 or GRA24 from the parasitophorous vacuole. **A**. Strategy for generating a disruption in *ROP17.* Plasmid pTKO2 containing HXGPRT (conferring resistance to mycophenolic acid in an otherwise Δ*hxgprt* strain) was integrated into a cleavage site generated by Cas9 in the beginning of the ROP17 open reading frame. Positions of the primers F and R that were used for detection of the insertion are shown. **B**. PCR data showing results of amplification with primers F and R of panel A. The wild-type locus yields a band of ~2275 bp whereas insertion of the knock-out plasmid yields a band of ~6000 bp. **C**. IFA of HFFs infected with RH-WT or RHΔ*rop17* that had also been transiently transfected with GRA16HA (left) or GRA24MYC (right) and then stained with anti-HA (red, left), or anti-Myc tag (red, right) and DAPI (blue) to reveal the nuclei. Hollow arrows indicate parasitophorous vacuoles; solid arrows indicate the nuclei in such cells. Translocation of GRA16 and GRA24 to the host nucleus is seen in cells infected with RH-WT but not RHΔ*rop17* parasites (quantitation of similar such experiments is shown in Fig. 5).

To confirm the importance of ROP17, we generated a complemented Δ*rop17* mutant in which a triple HA-tagged version of ROP17 is expressed off an introduced transgene (Fig. 3A). Within the parasite, ROP17HA colocalized with ROP2/3/4 at the apical end of the parasite (Fig. 3B), supporting proper localization of the tagged protein. As our previous results suggested ROP17 may be playing a critical role in the export of MYR-dependent proteins, we used the host c-Myc regulation phenotype as a readout of successful complementation. The results (Fig. 3C) showed that cells infected with the Δ*rop17* mutants show only background levels of c-Myc in the nuclei of infected cells whereas cells infected with wild type and the complemented mutant show robust c-Myc expression (confirmation and quantification of these results are presented further below). These results thereby confirm that ROP17 is indeed necessary for host c-Myc upregulation by *Toxoplasma* tachyzoites, and in combination with the defect of translocation of GRA16 and GRA24 this strongly suggests that ROP17 is a previously unknown player in the process whereby GRA proteins cross the PVM.

**Figure 3.**
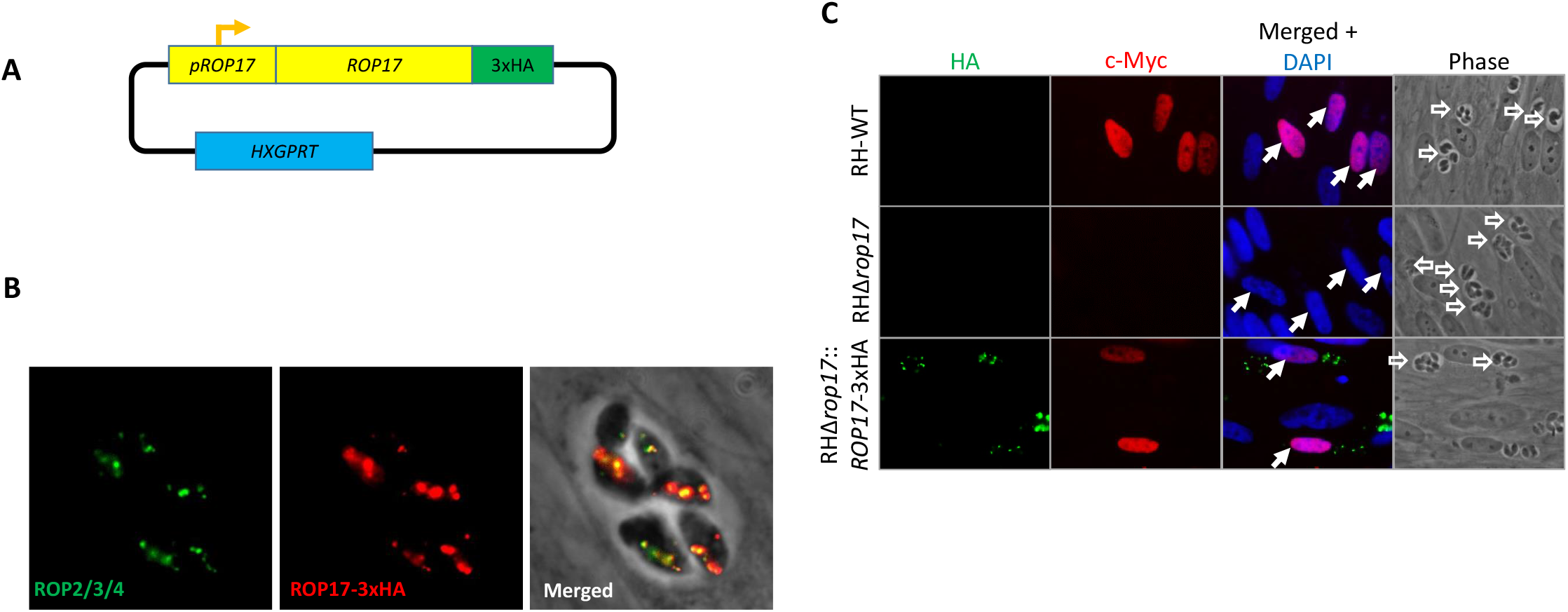
A wild-type copy of *ROP17* rescues the Myr^-^ phenotype of the Δ*rop17* mutant. **A**. Strategy for complementing the Δ*rop17* mutants with a 3xHA-tagged wild type copy of *ROP17.* **B**. IFA showing successful complementation of the Δ*rop17* mutant with a HA-tagged wild-type copy of the gene. Green shows staining with anti-ROP2/3/4 as a marker for rhoptries; red shows staining for the complementing ROP17-HA. **C**. IFA of HFFs infected with RH-WT, RHΔ*rop17* or RHΔ*rop17::ROP17-3xHA*. Anti-HA antibody was used to detect the complementing ROP17 (green) while red was used for staining of host c-Myc as an indicator of successful effector translocation, and blue shows DAPI staining of the host nuclei. Hollow arrows indicate parasitophorous vacuoles; solid arrows indicate the host nuclei in infected cells (quantitation of similar such experiments is shown in Fig. 5).

Even though ROP17 is a well-studied serine/threonine protein kinase, it is possible that its role in protein translocation at the PVM is as a scaffolding protein rather than as an active kinase. To test this, we made three different versions of ROP17, each with a mutation to alanine in one of three residues known to be essential for catalysis [44–46], i.e., K312A, D436A and D454A (Fig. 4A). These mutated versions were introduced into the Δ*rop17* mutant where they showed the expected colocalization with ROP2/3/4 in puncta at the apical end (Figure 4B). To determine if the mutant ROP17s reached the PVM after invasion, we applied previously established conditions for partially permeabilizing infected host cells such that antibodies can reach only the PVM, not the parasites within [47]. This showed that, indeed, in cells where the control antibody (anti-SAG1) fails to detect the parasites within the PVM, anti-ROP17 efficiently stains the PVM showing that the mutant ROP17s do successfully enter the host cell and traffic to this location (Fig. 4C).

**Figure 4.**
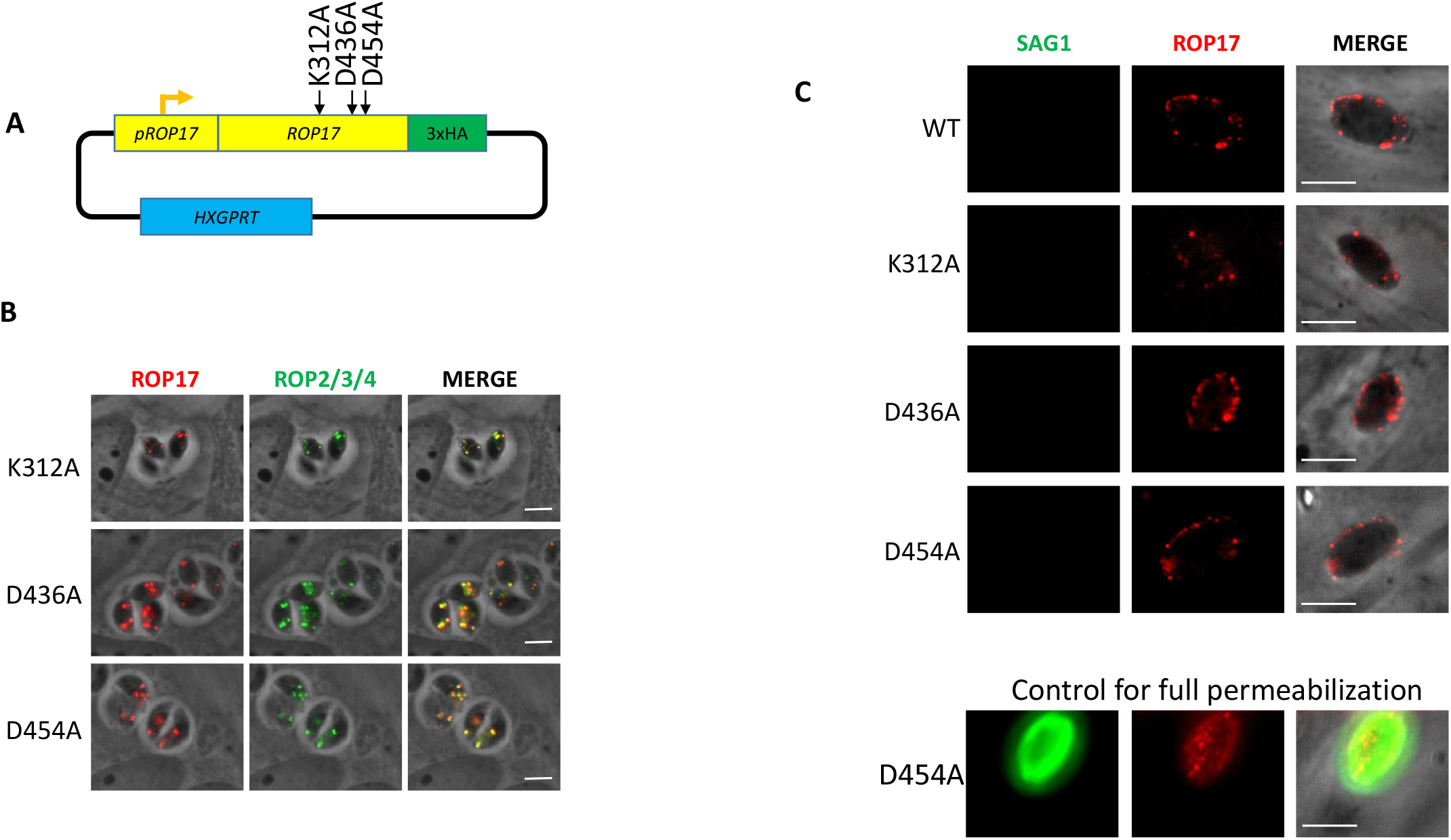
Creation of three strains of RHΔ*rop17* containing point mutations in key catalytic residues. **A**. Three different plasmids were created for the expression of point mutant variants of ROP17; each expresses a *ROP17* transgene encoding an alanine substitution at one of the three predicted catalytic residues: K312, D436 and D454. **B**. IFA of infected HFFs showing correct trafficking of the mutated ROP17 expressed in an RHΔ*rop17* background. Anti-HA (red) detects the ROP17 transgene product while anti-ROP2/3/4 (green) detects other known rhoptry proteins. **C**. IFA of HFFs that were infected with the strains expressing the indicated HA-tagged version of ROP17 (WT, K312A, D436A or D454A), partially permeabilized by treatment with 0.02% digitonin, and then stained for ROP17-3xHA using anti-HA (red) or anti-SAG1 (green). The absence of SAG1 staining was used to indicate that the parasitophorous vacuole was not permeabilized indicating partial permeabilization that allows antibodies to access the host cytosol but not penetrate the PVM. ROP17 is detected at the PVM in cells that are only partially permeabilized indicating the expected cytosolic exposure of the protein. A positive control of an infected cell that was fully permeabilized under these conditions is shown to confirm the anti-SAG1 staining is readily seen in such cells.

We next sought to determine if the ectopic expression of the wild type ROP17 gene and/or the mutant versions could complement the phenotypes we have observed in the ROP17-disrupted strains. When we assessed protein translocation of GRA24MYC, we observed translocation into the nucleus of ~89% of cells infected with GRA24MYC-expressing wild type parasites whereas cells infected with GRA24MYC-expressing RHΔ*rop17* parasites showed no such translocation (Fig. 5A). The loss of GRA24 translocation was successfully rescued when the RHΔ*rop17* parasites were complemented with a fully functional ROP17 but not with any of the point mutant versions of ROP17 (Fig. 5A). When we assessed c-Myc upregulation, a host phenotype associated with the translocation of MYR1-dependent effectors, we observed a similar result; c-Myc was upregulated in >90% of the nuclei of host cells infected with wild type RH and RHΔ*rop17::ROP17HA* parasites, but in <20% of the nuclei of host cells infected with RHΔ*rop17* or any of the three versions complemented with a mutation altering the trio of catalytic residues (Fig 5B, C). These results strongly suggest that ROP17’s kinase activity is necessary for its role in GRA translocation across the PVM.

**Figure 5.**
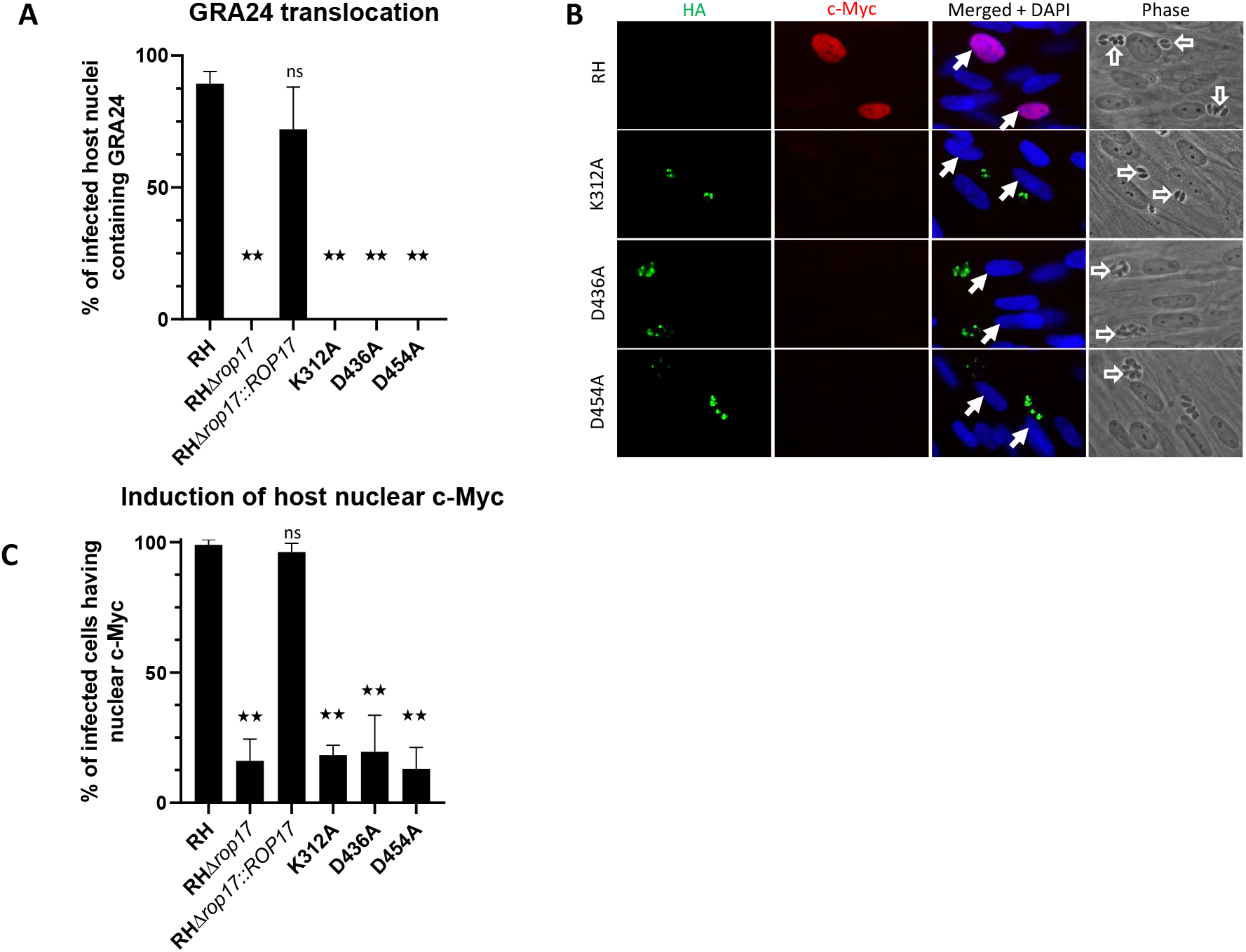
ROP17 catalytic activity appears necessary for its role in effector translocation. **A**. HFFs were infected with the indicated strains transiently expressing GRA24MYC and then 17-18 hours later the presence of GRA24MYC in the nucleus of cells infected with GRA24MYC-expressing parasites was assessed by IFA. The results are from assessment of a minimum of 94 cells and the standard error of the mean is shown. The experiment was done in biological duplicate for RH, RHΔ*rop17* and *RHΔrop17::ROP17* and similar results were obtained in both. Catalytically inactive mutants were done in technical triplicate. **B**. Assessment of the effect of mutating ROP17 on host c-Myc upregulation upon infection. HFFs were infected with the indicated strain and then 20 hours later, c-Myc expression in the nucleus of infected cells was assessed using IFA and anti-c-Myc antibodies (red). Expression of the variants of ROP17-3xHA was assessed by staining with anti-HA (green). None of the three catalytic mutants rescued the Myr-phenotype of the Δ*rop17* mutant. Differences were assessed by ANOVA with a post hoc Tukey’s test. **: p<0.0001. **C**. Quantification of cells upregulating host c-Myc. At least 110 random fields were quantified from the slides used to produce the images shown in panel B, scoring for percentage of host nuclei in infected cells that show host c-Myc upregulation. Differences were assessed by ANOVA with a post hoc Tukey’s test. **: p<0.0001.

Given that ROP17 ends up at the PVM in infected cells and given that this is where the known GRA translocation machinery (e.g., MYR1 /2/3) is located, it seemed most likely that this is where ROP17 functions to assist in the translocation of GRA proteins. To test this directly, we took advantage of the fact that when a tachyzoite infects a cell, the ROP proteins that are injected can associate either with the PVM of that parasite or with the PVM surrounding other parasites that are also present within that cell [48]. Thus, we created a reporter parasite line that lacked ROP17 expression but was stably expressing GRA24MYC; translocation of GRA24MYC in this strain should be blocked at the PV unless ROP17 can be provided in *trans.* We then infected cultures with these RHΔ*rop17::GRA24MYC* parasites, followed an hour later with RHΔ*myr1* parasites expressing mCherry to distinguish them from the nonfluorescent RHΔ*rop17::GRA24MYC* line (“Condition 1”, Fig. 6A). In case the order of infection was important, we also did this experiment where we inverted the order that the two strains were added to the monolayers; i.e., we initiated the infections with the RHΔ*myr1* mCherry line, followed an hour later by infection with the nonfluorescent *RHΔrop17::GRA24MYC* (“Condition 2,” Figure 6A). In both cases, we assessed GRA24MYC translocation in infected cells after a further 17 hours, scoring cells that were infected with either of the mutants alone or those co-infected with both. The prediction was that if ROP17’s role in GRA translocation is on the host-cytosolic side of the PVM, then the Δ*myr1* line would provide a functional ROP17 that could act in *trans* on the translocation machinery expressed by the Δ*rop17* parasites, whereas cells infected with either strain alone would not exhibit translocation. As shown in Figure 6B, this was indeed the result obtained; cells infected with either mutant alone showed no GRA24MYC in the host nucleus, whereas ~20-22% of co-infected cells did show translocation, regardless of the order that the two strains were added to the cultures. These results strongly argue that the action of ROP17 can be provided in *trans* and is needed within the host cytosol, not within the parasites or within the PV space since proteins are not known to be able to traffic across the PVM from host to PV or parasite (except to the lysosome for digestion [49]).

**Figure 6.**
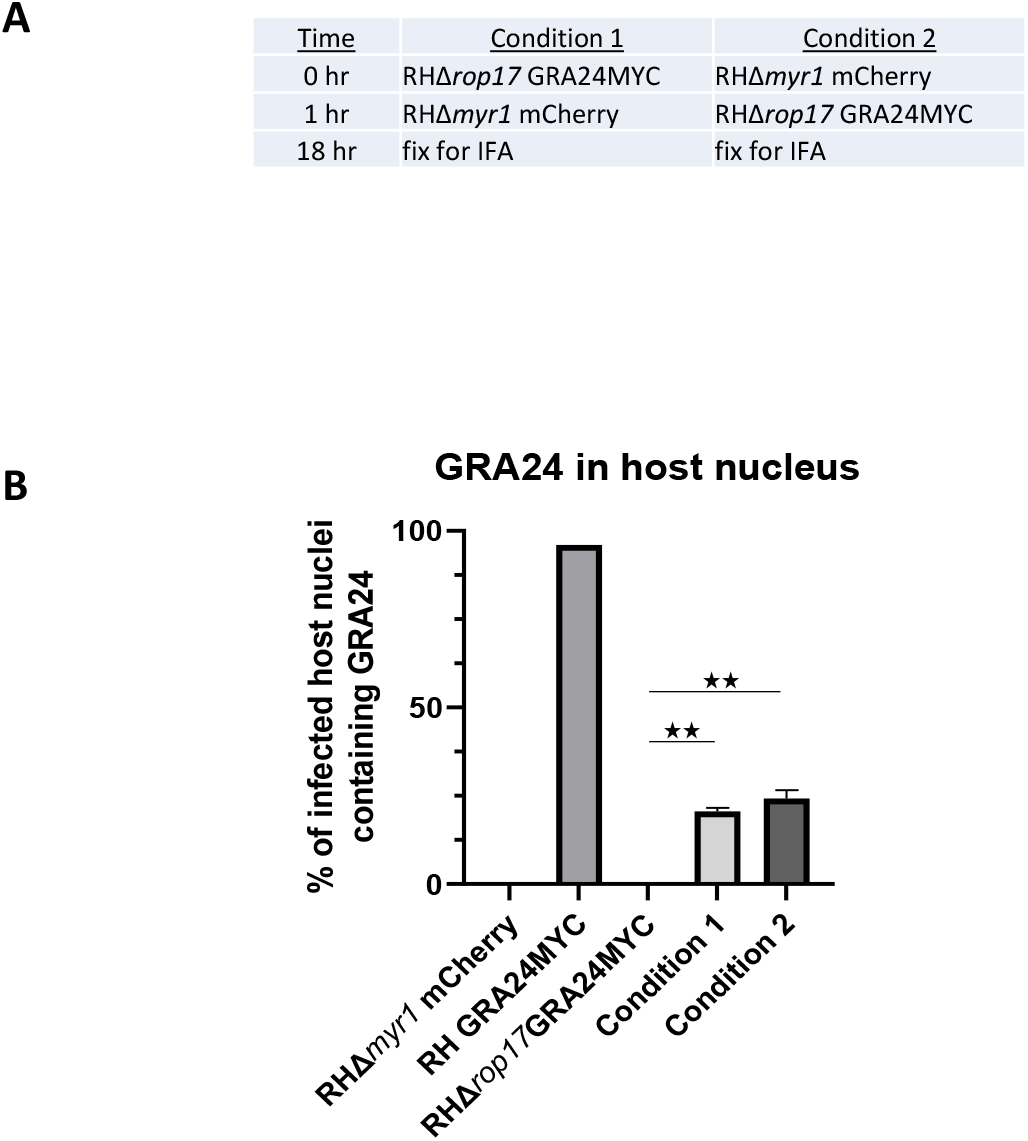
ROP17’s role in effector translocation occurs at the host-cytosolic side of the PVM. **A**. Conditions used for co-infection of host cells with RHΔ*rop17::GRA24MYC* and RHΔ*myr1* expressing mCherry. Infections were initiated with the indicated strain followed by addition of the second strain one hour later followed by IFA after a further 17 hours. **B**. Quantitation of the percentage of host nuclei staining positive for GRA24MYC in the cells infected with the indicated strains. Cells infected with one or other of the two mutants showed no GRA24MYC whereas those that were coinfected with both mutants showed substantial rescue in “trans.” Differences were assessed by ANOVA with a post hoc Tukey’s test. **: p<0.001.

The data presented so far show that ROP17 is necessary for the translocation of at least two GRA proteins from the PV to the host nucleus. To determine if this is true of essentially all dense granule effectors that end up in the host cell, we performed RNASeq analysis on HFFs infected with RH wild type vs. RHΔ*rop17* parasites at 6 hours post infection. As a control for a strain that has previously been shown by RNASeq analysis to be defective in the translocation of seemingly all soluble GRA effectors [41], we used a Δ*myr1* strain. The results showed substantial concordance between the genes modulated in a MYR1-dependent manner and a ROP17-dependent manner. This conclusion can be illustrated by Principal Component Analysis (PCA) of the host genes sets generated during infection with these strains. In Figure 7A, RNASeq data for the 8 strains analyzed are shown on the plot of the first and second principal component, and the individual genes along with their RPKM values are displayed in Supplemental Table S1. The samples infected with RHΔ*myr1* and RHΔ*rop17* cluster together closely and well apart from both mock-infected cells and RH-infected cells.

**Figure 7.**
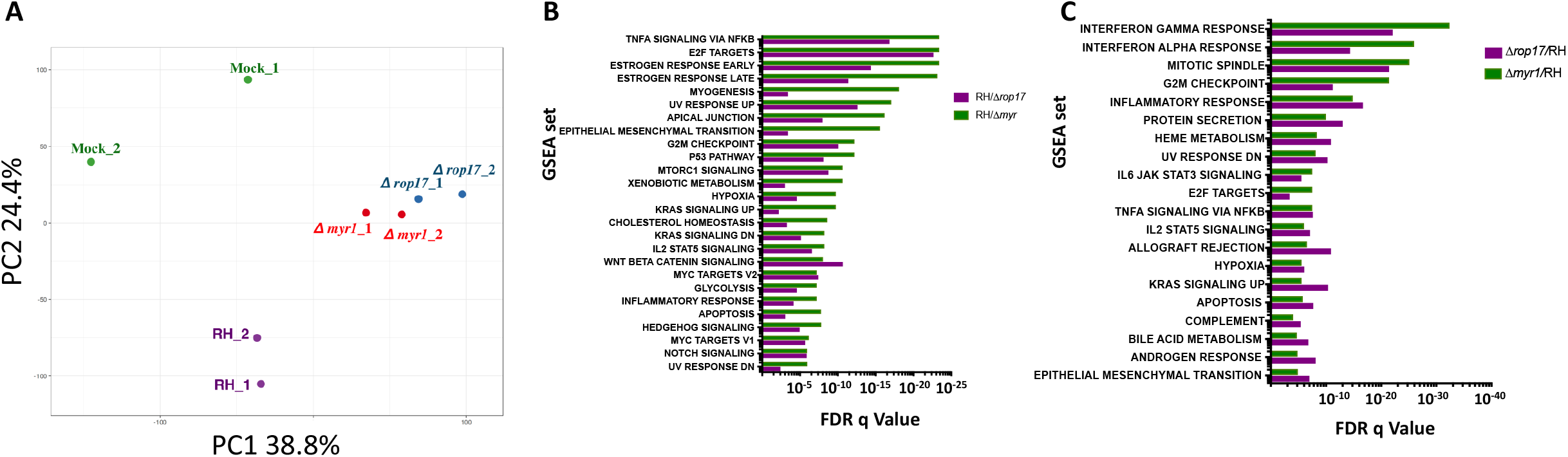
Disrupting *MYR1* or *ROP17* have congruent impacts on the infected host cell’s transcriptome as assessed by RNASeq. **A**. Principal component analysis (PCA) of the RPKM values of HFFs mock-infected or infected with RH-WT (wild type), RHΔ*myr*!, or RHΔ*rop17* tachyzoites. On a plot of PC1 and PC2, there is close similarity of the data for cells infected with the two mutants, RHΔ*myr1* and RHΔ*rop17*, relative to the mock- or RH-WT-infected cells. Gene names and RPKMs can be found in Supplemental table S1. **B**. Gene-set expression analysis (GSEA) of all genes expressed 2.5-fold higher in RH than in RHΔ*myr1* (green) or RHΔ*rop17* (purple), where either was lower than the FDR q-value threshold of 10^-5^. The gene sets are ordered based on descending levels of significance (ascending q-values) for the cells infected with RHΔ*myr1*. **C**. As in B, except GSEA was performed on the genes that were expressed 2.5-fold lower in RH compared to RHΔ*myr1* and RHΔ*rop17* by the same criteria.

To further explore this similarity, genes that exhibit a 2.5-fold difference during infection with these mutants compared to infection with the wild type were grouped by Gene Set Enrichment Analysis (GSEA). First, genes that were *increased* in cells infected with the wild type parasite (RH) compared to the two mutants were analyzed by GSEA. Figure 7B shows a list of gene sets in which the FDR q-value of either the RH vs. RHΔ*myr1* or the RH vs. RHΔ*rop17* was less than 10^-5^. For each of these gene sets, both the FDR q-value of the RH vs. RHΔ*myr1* (green) and RH vs. RHΔ*rop17* (purple) is shown. Genes that were expressed 2.5-fold *lower* in cells infected with the wild type compared to either of the mutants were also analyzed by GSEA and gene sets in which either the

FDR q-value of RH vs. RHΔ*myr1* (green) or RH vs. RHΔ*rop17* was less than 10^-5^ were plotted in Figure 7C. In both cases, almost all the gene sets that are strongly affected by the lack of *MYR1* were similarly affected by the lack of *ROP17,* although the magnitude of the effect varied somewhat. Finally, we performed a direct comparison of expression levels in cells infected with RHΔ*myr1* and cells infected with RHΔ*rop17* (instead of comparing each to the wild type-infected cells). We used GSEA to analyze genes for which expression was 2.5 fold higher or 2.5-fold lower in RHΔrop17-infected cells than in RHΔ*myr1*-infected cells. The results were that in neither case was a gene set enriched with an FDR q-value of even a very relaxed threshold of 10^-4^. Hence, the absence of ROP17 and MYR1 appear to have similar impacts on the infected cell and thus ROP17 appears necessary for the action of most, probably all, GRA effectors that transit across the PVM via the MYR machinery.

## Discussion

Using a genetic screen, we have identified the rhoptry-derived serine-threonine protein kinase, ROP17, as required for action of most if not all GRA effectors that translocate across the PVM. Using a cellular “trans” complementation assay, we have further shown that the role of ROP17 is within the host cytosol, not within the parasite or PV space, and that ROP17 must be catalytically active to accomplish this role. Given its location at the PVM [13,21], where other necessary elements of the translocation machinery are present, these results strongly argue that ROP17 acts on one or other components of this machinery on the host cytosol side. Although we cannot formally exclude the possibility that ROP17 assists GRA effectors in their trafficking across the host cytosol, from the PVM to the host nucleus, the fact that the GRA effectors that reach the host nucleus possess a conventional nuclear-localization signal (NLS) argues against this possibility as there should be no need for any additional help in their journey. Indeed, heterologous expression of GRA16 and GRA24 in uninfected cells shows results in efficient trafficking to the host nucleus, confirming that no parasite proteins are necessary for this last stage of their journey [36,37].

Given that ROP17 is a protein kinase, it seems most likely that its role in translocating GRA effectors is through phosphorylation of one or more key components of the translocation machinery. Phosphoproteomic analyses on cells infected with *Toxoplasma* tachyzoites revealed that many parasite proteins are phosphorylated at serine and threonine residues after their secretion from the parasite [50]. Among such proteins are the PVM-localized MYR1 and MYR3 that are known to be required for GRA translocation [42,43]. The protein kinases that mediate these phosphorylations have not yet been identified but protein phosphorylation is a well-established way to regulate protein function and so such modifications might be required for PVM-localized proteins like these to become activated for their respective roles. Efforts to determine the full machinery involved in GRA translocation across the PVM are underway and once the full complement of proteins is known, mapping of all their phosphosites and determination of which such sites are dependent on which protein kinase (e.g., ROP17 or, perhaps, ROP18, another serine-threonine kinase present at the PVM) and which of these sites must be phosphorylated for functional translocation will be an important follow-up to the work proposed here.

Both ROP17 and ROP18 are involved in the inactivation of immunity-related GTPases [15,16,21]. Our finding that ROP17 has at least two biological roles is similar to what has been reported for ROP18; this related ROP2-family member is involved in IRG inactivation and proteasomal degradation of ATF6beta, a host transcription factor that localizes to the host endoplasmic reticulum (which itself is adjacent to and maybe even contiguous with the PVM [51]) and that is crucial to the host immune response [52]. Interestingly, it is the N-terminal region of ROP18, which lies outside the conserved kinase domain, that binds to ATF6beta but an active kinase domain is required for the inactivation, suggesting that ATF6beta is a substrate for phosphorylation by ROP18 [52]. Our results add a further possible explanation for the previously reported attenuation of ROP17 mutants in a mouse model of virulence using a Type I strain [26]; i.e., the decrease in virulence could be due to some combination of a weakened defense against IRGs and the defect in GRA effector translocation reported here. A major role for the latter would be consistent with the previously reported attenuation in Type I Δ*myr1* strains in a similar mouse model [42].

The involvement of a rhoptry protein in the function of GRA proteins is a second example of “inter-organellar” collaboration, the first being the binding of micronemal AMA1 to rhoptry neck protein, RON2, during the invasion of tachyzoites into the host cell [53,54]. This second example wherein a rhoptry bulb protein, ROP17, somehow assists in the translocation of GRA proteins makes clear that these different secretory organelles are part of a complicated but concerted machinery used by the parasites to interact with the host cell they are infecting.

Finally, it is worth noting that the chemical mutagenesis used to generate the mutant library that yielded these *ROP17* mutants provided more information than just the fact that this protein plays an important role. By specifically looking for hypomorph mutants that showed only a partial defect we were able to identify a missense M350K mutation in ROP17 providing structure/function information, namely that this residue is important for this function of ROP17. This site is within a predicted loop region that is just N-terminal of beta-sheet 4 and well away from the active site [13,44,45]. Interestingly, this is part of a region that is highly variable between different members of the ROP2 family but is in a stretch of about 9 residues that is not present in other protein kinases [45]. This may be related to the unusual, multiple functions of these secreted kinases, perhaps in enabling them to target specific substrates at this crucial interface of host and parasite.

## Materials and Methods

### Parasite culture

*Toxoplasma gondii RHΔhpt* [55] was used for this study. *Toxoplasma* tachyzoites were maintained by serial passage in human foreskin fibroblasts (HFFs) cultured in complete Dulbecco’s Modified Eagle Medium (cDMEM) supplemented with 10% heat-inactivated fetal bovine serum (FBS), 2 mM L-glutamine, 100 U/ml penicillin and 100 μg/ml streptomycin and grown at 37 °C in 5% CO2. Infections included in this study were performed by scraping infected monolayers and lysing the host cells open using a 27-gauge needle. The released parasites were pelleted at 1500 rpm for 10 min, resuspended, counted using a hemocytometer, and added to confluent HFFs at the multiplicity of infection (MOI) stated.

### Genome sequencing

For whole genome sequencing on the parental RH strain (SRR2068658), MFM1.15 (SRR5643318), and MFM2.1b (SRS2249312) mutants, a single Illumina PE barcoded library was prepared from tachyzoite gDNA. Libraries were then pooled into groups of nine samples and multiplex sequenced in a single lane of an Illumina HiSeq 2000 machine to generate about 3 Gb of sequencing data per sample. As the mutants were made using the Type I RH strain as the parent, the sequencing reads were first quality trimmed with trimommatic and then mapped to the reference assembly of the Type I GT1 strain (as present in ToxoDB v13.0) with *bowtie2* [56]. After removing duplicated reads with *picard* and adjusting alignments around indels with *GATK toolkit* [57], single nucleotide variants (SNVs) were called using samtools utility *mpileup* [58] requiring a minimum base coverage of 5 reads and an alternative allele frequency of at least 80% or higher. Following this, *SnpEff* [59] together with a gff3 annotation file from the reference GT1 strain (ToxoDB v13.0) were used to classify the different types of SNVs present in each mutant. Potential change-of-function SNVs that were different between any of the two mutants and both the parental and reference strains were selected for further quality control and analysis.

### Transfections

All transfections were performed using the *BTX* EMC600 *Electroporation System* (Harvard Apparatus) or Amaxa 4D Nucleofector (Lonza) model. Tachyzoites were mechanically released in PBS, pelleted, and resuspended in solution for transfection. After transfection, parasites were allowed to infect HFFs in DMEM. Transfections with the *BTX* EMC600 model were performed using 5-10 x 10^6^ parasites and 5-10 μg DNA in Cytomix (10 mM KPO_4_, pH 7.6, 120 mM KCl, 5 mM MgCl_2_, 25 mM HEPES, 2 mM EDTA, 150 μM CaCl_2_). Transfections with the Amaxa 4D model were performed using 1-2 x 10^6^ parasites in 20 μl P3 solution or 5-10 x 10^6^ parasites in 100 μl P3 solution with 5-15 μg DNA. Effector translocation assays were performed by transiently transfecting pHTU-GRA24MYC [43] or pGRA1-GRA16HA [43] plasmid into tachyzoites, infecting monolayers of HFFs in DMEM, and fixing monolayers with formaldehyde at 16-24 hpi.

### Immunofluorescence microscopy

Infected cells grown on glass coverslips were fixed using methanol at −20 °C for 20 min or 4% formaldehyde at room temperature (RT) for 20 min, as stated in the text. Methanol-fixed samples were washed three times for 5 min with PBS and blocked using 3% BSA in PBS for 1 hr at RT. Formaldehyde-fixed samples were rinsed once with PBS, permeabilized with 0.2% Triton-X 100 (TTX-100) for 20 min, and then blocked as described above. GRA16HA (and other HA-tagged proteins) was detected using rat anti-HA antibodies (Roche) while GRA24MYC was detected using rabbit anti-MYC tag antibody 9E10 (Santa Cruz Biotechnology). This anti-MYC tag antibody does not detect host c-Myc. Host c-Myc was detected using monoclonal antibody Y69, which does not cross react with the MYC tag expressed by GRA24MYC. Primary antibodies were detected with goat polyclonal Alexa Fluor-conjugated secondary antibodies (Invitrogen). Vectashield with DAPI stain (Vector Laboratories) was used to mount the coverslips on slides. Fluorescence was detected using a LSM710 inverted confocal microscope (Zeiss) or epifluorescence microscope, as stated in the text. Images were analyzed using ImageJ. All images shown for any given condition/staining in any given comparison/dataset were obtained using identical parameters.

### Quantitation of nuclear GRA24MYC

To assess the amount of GRA24MYC that translocated to the nucleus following transient transfection, phase contrast, DAPI and anti-MYC tag images were taken of 10-20 fields of view containing tachyzoites-infected HFF at 20 hours PI. Phase contrast was used to define the infected cells of these images, then ImageJ was used to define the nucleus on the DAPI-stained corresponding images, and these nuclear boundaries were then quantified for the intensity of GRA24MYC intensity on the corresponding MYC-tag stained images.

### Partial Permeabilization

Parasites were syringe-released using a 27g needle and used to infect HFFs for 2 hrs, at which time the cells were washed with PBS and then fixed with 4% formaldehyde at room temperature (RT) for 20 min. Formaldehyde-fixed samples were rinsed once with PBS, permeabilized with 0.02% digitonin solution for 5 min and then blocked with 3% BSA in PBS for 1 hr at RT. Staining was performed with anti-HA (Roche) and anti-SAG1 (DG52) primary antibodies and polyclonal Alexa Fluor-conjugated secondary antibodies (Invitrogen). Partial permeabilization of a particular vacuole was determined by the exclusion of the SAG1 antibody.

### Gene disruption

The RHΔ*rop17* strain was generated by disrupting the corresponding gene locus using CRISPR-Cas9 and selecting for integration of a linearized vector encoding hypoxanthine-guanine phosphoribosyl transferase *(HXGPRT)* using drug selection for 8 days using 25 μg/mL mycophenolic acid (MPA) and 50 μg/mL xanthine (XAN) for HXGPRT selection. Specifically, the pSAG1:U6-Cas9:sgUPRT vector [60] was modified by Q5 site-directed mutagenesis (NEB) to specify sgRNAs targeting *ROP17* (F2). The resulting sgRNA plasmid, dubbed pSAG1:U6-Cas9:sgROP17 (P1) was transfected into the RHΔhpt strain of *Toxoplasma* with pTKO2 (HXGPRT+) plasmid. The parasites were allowed to infect HFFs in 24-well plates for 24 hrs, after which the media was changed to complete DMEM supplemented with 50 μg/ml mycophenolic acid (MPA) and 50 μg/ml xanthine (XAN) for HXGPRT selection. The parasites were passed twice before being single cloned into 96-well plates by limiting dilution. Disruption of the gene coding regions was confirmed by PCR and sequencing of the locus.

### Ectopic gene integration

The RHΔ*rop17* strain was complemented ectopically with the pGRA-ROP17-3xHA plasmid, which expresses *ROP17* off its natural promoter. To construct the pGRA-ROP17-3xHA plasmid, pGRA1plus-HPT-3xHA plasmid [43] was first digested using EcoRV-HF and NcoI (New England Biolabs) for 4 hrs at 37 °C to remove the *GRA1* promoter. Product was incubated with Antarctic phosphatase (New England Biolabs) and gel-extracted. The empty vector backbone was amplified by PCR using Herculase II polymerase (Agilent) and primers 5’-CACATTTGTGTCACCCCAAATGAGAATTCGATATCAAGCTTGATCAGCAC-3’ and 5’-GAGGCGGCTTTATTACAGAAGGAGCCATGGTACCCGTACGACGTCCCG-3’ with each having 23 and 24 base pair overhangs to *ROP17,* respectively. The *ROP17* promoter and open reading frame were amplified from RHΔhpt genomic DNA and using 5’-GTGCTGATCAAGCTTGATATCGAATTCTCATTTGGGGTGACACAAATGTG-3’ and 5’-CGGGACGTCGTACGGGTACCATGGCTCCTTCTGTAATAAAGCCGCCTC-3’ primers, each containing 27 or 24 base pair overhangs to the pGRA1plus-HPT-3xHA plasmid backbone, respectively. Amplified backbone and *ROP17* were then assembled using the Gibson assembly master mix (New England Biolabs). ElectroMAX DH10B *E. coli* (Invitrogen) were subsequently transformed and plated to obtain single colonies of successfully assembled pGRA1-ROP17-3xHA plasmid. ROP17 integration was verified by PCR and sequencing using primers 5’-CACTGATCGGCTTTGTAGACTT-3’ and 5’-CGCGCACGGCAGTCAGATAA-3’.

To complement RHΔ*rop17Δhpt* parasites with wildtype *ROP17,* the pGRA1-ROP17-3xHA plasmid construct described previously was transfected to generate an RHΔ*rop17::ROP17* population. This population was selected by MPA/XAN as previously described. The resulting population was then cloned by limiting dilution and tested for ROP17-3xHA expression by Western blot and IFA.

To generate RH and RHΔ*rop17* parasite lines ectopically expressing GRA24MYC, pHTU-GRA24-3xMYC was transfected into each strain and selected using MPA/XAN as described above for 6 days.

### Site-specific mutation

Site-specific point mutation of the pGRA-ROP17-3xHA plasmid was performed by creating primers 5’-GCGATATTTGTTCAACGGGTGTTGAGCAAT-3’ and 5’-CAGCGCGAATGGTTGCCCTGTGGTGGG-3’, 5’-GCTGTGAAACTGCAAAATTTTCTTGTTGAT-3’ and 5’-GCCATGAACAAGTCCGAACGCGTGGAA-3’, and 5’-GcCTTCACTCAAATTCTTCGTACGAATG-3’ and 5’-AGAAAGTAGAAGCAATCCCGATTTATC-3’ to mutate residues 312, 436, and 454, respectively, to an alanine codon within the *ROP17* open reading frame. The “Round-the-horn” site-directed mutagenesis approach was used to introduce point mutations at the aforementioned residues using the ROP17-3xHA plasmid. The PCR products were then individually ligated using a KLD Enzyme reaction kit (New England Biolabs) for 3 hours and subsequently transformed into ElectroMAX DH10B *E. coli* (Invitrogen). Single colonies for each point mutant were Miniprepped (Qiagen) and sequence-verified using either 5’-GCCATGAACAAGTCCGAACGCGTGGAA −3’ or 5’-GCGATATTTGTTCAACGGGTGTTGAGCAAT −3’.

To generate parasite lines complemented with the catalytically inactive *ROP17,* RHΔ*rop17* parasites were transfected with the pGRA1-ROP17_K312A_-3xHA, pGRA1-ROP 17_D436A_-3xHA, or pGRA1-ROP17_D454A_-3xHA plasmid then subsequently selected for 6 days with MPA/XAN and single cloned as previously described.

### Coinfection Assays

Condition 1. Confluent HFF coverslips were infected with RHΔ*rop17* parasites stably expressing GRA24-3xMYC at an MOI of 0.15. They were then pulsed at 1400 rpm and placed at 37°C and 5% CO2 for 1 hour. Thereafter, the same sample was infected with RHΔ*myr1* constitutively expressing mCherry at a MOI of 0.15, pulsed at 1400 rpm and placed at 37°C and 5% CO2. Infections were allowed to progress to 17-18 hours. Condition 2 was performed in the same way except the order of addition of the two strains to the host cells was reversed.

### RNA extraction, library preparation, and sequencing

HFFs were serum-starved for 24 hours before infection by growth in DMEM containing 0.5% serum. They were then infected with the indicated line of tachyzoites at an MOI of 5, and at 6 hpi, 1 ml TRIzol reagent (Invitrogen) was added to each T25 and the cells were scraped. Lysates were collected and frozen at −20 °C. Total RNA was extracted following the manufacturer’s instructions, with some modifications. Briefly: frozen samples were thawed on ice and 0.2 ml chloroform was added to TRIzol suspensions, which were then mixed by inverting 10 times, and incubated for 5 min. Tubes were then spun at 12,000 rpm for 15 min at 4 °C. RNA in the aqueous phase was transferred into a fresh tube and 0.5 ml absolute isopropyl alcohol was added and incubated at 4 °C for 10 min. They were then spun at 12,000 rpm for 20 min at 4 °C. After decanting the supernatants, RNA pellets were washed with 1 ml 75% ethanol and then spun at 12,000 rpm for 20 min at 4 °C. Supernatants were removed and the RNA pellets were resuspended in 30 μl RNase-free DEPC-water. RNA samples were submitted to the Stanford University Functional Genomic Facility (SFGF) for purity analysis using the Agilent 2100 Bioanalyzer. Multiplex sequencing libraries were generated with RNA Sample Prep Kit (Illumina) according to manufacturer’s instructions and pooled for a single high-throughput sequencing run using the Illumina NextSeq platform (Illumina Nextseq 500 model instrument).

### RNASeq read mapping and differential expression analysis

Raw reads were uploaded onto the CLC Genomics Workbench 8.0 (Qiagen) platform for independent alignments against the human genomes (Ensembl.org/ hg19) and *Toxoplasma* Type I GT1 strain (ToxoDB-24, GT1 genome). All parameters were left at their default values. The number of total reads mapped to each genome was used to determine the RPKM (Reads Per Kilobase of transcript per Million mapped reads). Among these genes, only those with an average RPKM ratios ≥2.5 were counted as changed in expression.

### Gene Set Enrichment Analysis (GSEA)

GSEA, available through the Broad Institute at http://www.broadinstitute.org/gsea/index.jsp, was the enrichment analysis software we used to determine whether defined sets of differentially expressed human genes in our experiment show statistically significant overlap with gene sets in the curated Molecular Signatures Databases (MsigDB) Hallmark gene set collection. We used the cutoff of FDR q-value <10^-5^.

### PCA Analysis

To generate PCA (principle component analysis) we used https://biit.cs.ut.ee/clustvis/#tab-9298-7 online tool. We used the RPKM values of all expressed genes to generate the PCA.

### Statistical Analyses

Statistical analysis was performed with Prism version 8 software. For intensity analysis, GRA24MYC translocation was assessed by ImageJ and then differences in intensity were analyzed by one way ANOVA with a post hoc Tukey’s test. Similarly, differences in the number of infected host cells with nuclei staining positive for GRA24 translocation or c-Myc expression were compared using a one way ANOVA with a post hoc Tukey’s test.

### Accession number

The RNASeq data files have been deposited in GEO under accession number GSExxxxx (number available at time of publish). Data presented as transcriptomics control data, for HFFs uninfected and those infected by RH and RHΔ*myr1* tachyzoites, have been published under GSE122786 (Panas et al., 2019, manuscript in press at mBio).

## Acknowledgments

We thank all members of our laboratory for helpful comments and input to the experiments and manuscript and Melanie Espiritu for help with tissue culture and ordering. We also thank Dr. Michael Reese (UT Southwestern) for help positioning the missense point mutation in the ROP17 secondary structure.

---

Supplemental Table S1. List of the RPKM values of HFF infected with the respective strains.

## References

1. Carruthers VB, Sibley LD (1997) Sequential protein secretion from three distinct organelles of *Toxoplasma gondii* accompanies invasion of human fibroblasts. Eur J Cell Biol 73: 114–123.

2. Hakansson S, Carruthers V, Heuser J, Sibley D (1997) A putative role for MIC2 in gliding motility of *Toxoplasma gondii*. Molecular Parasitology Meeting VIII, MBL Woods Hole, MA: 421.

3. Soldati D, Dubremetz JF, Lebrun M (2001) Microneme proteins: structural and functional requirements to promote adhesion and invasion by the apicomplexan parasite *Toxoplasma gondii*. Int J Parasitol 31: 1293–1302.

4. Boothroyd JC, Dubremetz JF (2008) Kiss and spit: the dual roles of *Toxoplasma* rhoptries. Nat Rev Microbiol 6: 79–88.

5. Tyler JS, Boothroyd JC (2011) The C-terminus of *Toxoplasma* RON2 provides the crucial link between AMA1 and the host-associated invasion complex. PLoS Pathog 7: e1001282.

6. Lamarque M, Besteiro S, Papoin J, Roques M, Vulliez-Le Normand B, et al. (2011) The RON2-AMA1 interaction is a critical step in moving junction-dependent invasion by apicomplexan parasites. PLoS Pathog 7: e1001276.

7. Tonkin ML, Roques M, Lamarque MH, Pugniere M, Douguet D, et al. (2011) Host cell invasion by apicomplexan parasites: insights from the co-structure of AMA1 with a RON2 peptide. Science 333: 463–467.

8. Kimata I, Tanabe K (1987) Secretion by *Toxoplasma gondii* of an antigen that appears to become associated with the parasitophorous vacuole membrane upon invasion of the host cell. J Cell Sci 88 (Pt 2): 231–239.

9. Dubremetz JF (2007) Rhoptries are major players in *Toxoplasma gondii* invasion and host cell interaction. Cell Microbiol 9: 841–848.

10. Bradley PJ, Sibley LD (2007) Rhoptries: an arsenal of secreted virulence factors. Curr Opin Microbiol 10: 582–587.

11. Hakimi MA, Olias P, Sibley LD (2017) *Toxoplasma* Effectors Targeting Host Signaling and Transcription. Clin Microbiol Rev 30: 615–645.

12. Kemp LE, Yamamoto M, Soldati-Favre D (2013) Subversion of host cellular functions by the apicomplexan parasites. FEMS Microbiol Rev 37: 607–631.

13. El Hajj H, Demey E, Poncet J, Lebrun M, Wu B, et al. (2006) The ROP2 family of *Toxoplasma gondii* rhoptry proteins: proteomic and genomic characterization and molecular modeling. Proteomics 6: 5773–5784.

14. Zhao Y, Ferguson DJ, Wilson DC, Howard JC, Sibley LD, et al. (2009) Virulent *Toxoplasma gondii* evade immunity-related GTPase-mediated parasite vacuole disruption within primed macrophages. J Immunol 182: 3775–3781.

15. Fentress SJ, Behnke MS, Dunay IR, Mashayekhi M, Rommereim LM, et al. (2010) Phosphorylation of immunity-related GTPases by a *Toxoplasma* gondii-secreted kinase promotes macrophage survival and virulence. Cell Host Microbe 8: 484–495.

16. Steinfeldt T, Konen-Waisman S, Tong L, Pawlowski N, Lamkemeyer T, et al. (2010) Phosphorylation of mouse immunity-related GTPase (IRG) resistance proteins is an evasion strategy for virulent *Toxoplasma gondii*. PLoS Biol 8: e1000576.

17. Howard JC, Hunn JP, Steinfeldt T (2011) The IRG protein-based resistance mechanism in mice and its relation to virulence in *Toxoplasma gondii*. Curr Opin Microbiol 14: 414–421.

18. Behnke MS, Fentress SJ, Mashayekhi M, Li LX, Taylor GA, et al. (2012) The polymorphic pseudokinase ROP5 controls virulence in *Toxoplasma gondii* by regulating the active kinase ROP18. PLoS Pathog 8: e1002992.

19. Fleckenstein MC, Reese ML, Konen-Waisman S, Boothroyd JC, Howard JC, et al. (2012) A *Toxoplasma gondii* pseudokinase inhibits host IRG resistance proteins. PLoS Biol 10: e1001358.

20. Niedelman W, Gold DA, Rosowski EE, Sprokholt JK, Lim D, et al. (2012) The rhoptry proteins ROP18 and ROP5 mediate *Toxoplasma gondii* evasion of the murine, but not the human, interferon-gamma response. PLoS Pathog 8: e1002784.

21. Etheridge RD, Alaganan A, Tang K, Lou HJ, Turk BE, et al. (2014) The *Toxoplasma* pseudokinase ROP5 forms complexes with ROP18 and ROP17 kinases that synergize to control acute virulence in mice. Cell Host Microbe 15: 537–550.

22. Sadak A, Taghy Z, Fortier B, Dubremetz JF (1988) Characterization of a family of rhoptry proteins of *Toxoplasma gondii*. Mol Biochem Parasitol 29: 203–211.

23. Beckers CJM, Dubremetz JF, Mercereau-Puijalon O, Joiner KA (1994) The *Toxoplasma gondii* rhoptry protein ROP2 is inserted into the parasitophorous vauole membrane, surrounding the intracellular parasite, and is exposed to the host cell cytoplasm. J Cell Biol 127: 947–961.

24. El Hajj H, Lebrun M, Fourmaux MN, Vial H, Dubremetz JF (2007) Inverted topology of the *Toxoplasma gondii* ROP5 rhoptry protein provides new insights into the association of the ROP2 protein family with the parasitophorous vacuole membrane. Cell Microbiol 9: 54–64.

25. Li JX, He JJ, Elsheikha HM, Chen D, Zhai BT, et al. (2019) *Toxoplasma gondii* ROP17 inhibits the innate immune response of HEK293T cells to promote its survival. Parasitol Res.

26. Fox BA, Sanders KL, Rommereim LM, Guevara RB, Bzik DJ (2016) Secretion of Rhoptry and Dense Granule Effector Proteins by Nonreplicating *Toxoplasma gondii* Uracil Auxotrophs Controls the Development of Antitumor Immunity. PLoS Genet 12: e1006189.

27. Cesbron-Delauw MF, Lecordier L, Mercier C (1996) Role of secretory dense granule organelles in the pathogenesis of *Toxoplasmosis*. Curr Top Microbiol Immunol 219: 59–65.

28. Charif H, Darcy F, Torpier G, Cesbron-Delauw MF, Capron A (1990) *Toxoplasma gondii:* characterization and localization of antigens secreted from tachyzoites. Exp Parasitol 71: 114–124.

29. Leriche MA, Dubremetz JF (1990) Exocytosis of *Toxoplasma gondii* dense granules into the parasitophorous vacuole after host cell invasion. Parasitol Res 76: 559–562.

30. Dubremetz JF, Achbarou A, Bermudes D, Joiner KA (1993) Kinetics and pattern of organelle exocytosis during *Toxoplasma gondii/host-cell* interaction. Parasitol Res 79: 402–408.

31. Mercier C, Cesbron-Delauw MF, Sibley LD (1998) The amphipathic alpha helices of the toxoplasma protein GRA2 mediate post-secretory membrane association. J Cell Sci 111: 2171–2180.

32. Mercier C, Howe DK, Mordue D, Lingnau M, Sibley LD (1998) Targeted disruption of the GRA2 locus in Toxoplasma gondii decreases acute virulence in mice. Infect Immun 66: 4176–4182.

33. Pernas L, Adomako-Ankomah Y, Shastri AJ, Ewald SE, Treeck M, et al. (2014) *Toxoplasma* effector MAF1 mediates recruitment of host mitochondria and impacts the host response. PLoS Biol 12: e1001845.

34. Rosowski EE, Lu D, Julien L, Rodda L, Gaiser RA, et al. (2011) Strain-specific activation of the NF-kappaB pathway by GRA15, a novel *Toxoplasma gondii* dense granule protein. J Exp Med 208: 195–212.

35. Hakimi MA, Bougdour A (2015) *Toxoplasma’s* ways of manipulating the host transcriptome via secreted effectors. Curr Opin Microbiol 26: 24–31.

36. Bougdour A, Durandau E, Brenier-Pinchart MP, Ortet P, Barakat M, et al. (2013) Host cell subversion by *Toxoplasma* GRA16, an exported dense granule protein that targets the host cell nucleus and alters gene expression. Cell Host Microbe 13: 489–500.

37. Bougdour A, Tardieux I, Hakimi MA (2014) *Toxoplasma* exports dense granule proteins beyond the vacuole to the host cell nucleus and rewires the host genome expression. Cell Microbiol 16: 334–343.

38. Gay G, Braun L, Brenier-Pinchart MP, Vollaire J, Josserand V, et al. (2016) *Toxoplasma gondii* TgIST co-opts host chromatin repressors dampening STAT1-dependent gene regulation and IFN-gamma-mediated host defenses. J Exp Med 213: 1779–1798.

39. Olias P, Etheridge RD, Zhang Y, Holtzman MJ, Sibley LD (2016) *Toxoplasma* Effector Recruits the Mi-2/NuRD Complex to Repress STAT1 Transcription and Block IFN-gamma-Dependent Gene Expression. Cell Host Microbe 20: 72–82.

40. He H, Brenier-Pinchart MP, Braun L, Kraut A, Touquet B, et al. (2018) Characterization of a *Toxoplasma* effector uncovers an alternative GSK3/beta-catenin-regulatory pathway of inflammation. Elife 7.

41. Naor A, Panas MW, Marino N, Coffey MJ, Tonkin CJ, et al. (2018) MYR1-Dependent Effectors Are the Major Drivers of a Host Cell’s Early Response to *Toxoplasma*, Including Counteracting MYR1-Independent Effects. MBio 9.

42. Franco M, Panas MW, Marino ND, Lee MC, Buchholz KR, et al. (2016) A Novel Secreted Protein, MYR1, Is Central to *Toxoplasma’s* Manipulation of Host Cells. MBio 7: e02231–02215.

43. Marino ND, Panas MW, Franco M, Theisen TC, Naor A, et al. (2018) Identification of a novel protein complex essential for effector translocation across the parasitophorous vacuole membrane of *Toxoplasma gondii*. PLoS Pathog 14: e1006828.

44. Qiu W, Wernimont A, Tang K, Taylor S, Lunin V, et al. (2009) Novel structural and regulatory features of rhoptry secretory kinases in *Toxoplasma gondii*. EMBO J 28: 969–979.

45. Labesse G, Gelin M, Bessin Y, Lebrun M, Papoin J, et al. (2009) ROP2 from *Toxoplasma gondii:* a virulence factor with a protein-kinase fold and no enzymatic activity. Structure 17: 139–146.

46. Peixoto L, Chen F, Harb OS, Davis PH, Beiting DP, et al. (2010) Integrative genomic approaches highlight a family of parasite-specific kinases that regulate host responses. Cell Host Microbe 8: 208–218.

47. Dunn JD, Ravindran S, Kim SK, Boothroyd JC (2008) The *Toxoplasma gondii* dense granule protein GRA7 is phosphorylated upon invasion and forms an unexpected association with the rhoptry proteins ROP2 and ROP4. Infect Immun 76: 5853–5861.

48. Hakansson S, Charron AJ, Sibley LD (2001) *Toxoplasma* evacuoles: a two-step process of secretion and fusion forms the parasitophorous vacuole. EMBO J 20: 3132–3144.

49. Dou Z, McGovern OL, Di Cristina M, Carruthers VB (2014) *Toxoplasma gondii* ingests and digests host cytosolic proteins. MBio 5: e01188–01114.

50. Treeck M, Sanders JL, Elias JE, Boothroyd JC (2011) The phosphoproteomes of *Plasmodium falciparum* and *Toxoplasma gondii* reveal unusual adaptations within and beyond the parasites’ boundaries. Cell Host Microbe 10: 410–419.

51. Goldszmid RS, Coppens I, Lev A, Caspar P, Mellman I, et al. (2009) Host ER-parasitophorous vacuole interaction provides a route of entry for antigen cross-presentation in *Toxoplasma* gondii-infected dendritic cells. J Exp Med 206: 399–410.

52. Yamamoto M, Ma JS, Mueller C, Kamiyama N, Saiga H, et al. (2011) ATF6beta is a host cellular target of the Toxoplasma gondii virulence factor ROP18. J Exp Med 208: 1533–1546.

53. Alexander DL, Mital J, Ward GE, Bradley P, Boothroyd JC (2005) Identification of the moving junction complex of *Toxoplasma gondii:* a collaboration between distinct secretory organelles. PLoS Pathog 1: e17.

54. Besteiro S, Michelin A, Poncet J, Dubremetz JF, Lebrun M (2009) Export of a *Toxoplasma gondii* rhoptry neck protein complex at the host cell membrane to form the moving junction during invasion. PLoS Pathog 5: e1000309.

55. Fox BA, Ristuccia JG, Gigley JP, Bzik DJ (2009) Efficient gene replacements in *Toxoplasma gondii* strains deficient for nonhomologous end joining. Eukaryot Cell 8: 520–529.

56. Langmead B, Salzberg SL (2012) Fast gapped-read alignment with Bowtie 2. Nat Methods 9: 357–359.

57. McKenna A, Hanna M, Banks E, Sivachenko A, Cibulskis K, et al. (2010) The Genome Analysis Toolkit: a MapReduce framework for analyzing next-generation DNA sequencing data. Genome Res 20: 1297–1303.

58. Li H, Handsaker B, Wysoker A, Fennell T, Ruan J, et al. (2009) The Sequence Alignment/Map format and SAMtools. Bioinformatics 25: 2078–2079.

59. Cingolani P, Platts A, Wang le L, Coon M, Nguyen T, et al. (2012) A program for annotating and predicting the effects of single nucleotide polymorphisms, SnpEff: SNPs in the genome of Drosophila melanogaster strain w1118; iso-2; iso-3. Fly (Austin) 6: 80–92.

60. Shen B, Brown KM, Lee TD, Sibley LD (2014) Efficient gene disruption in diverse strains of *Toxoplasma gondii* using CRISPR/CAS9. MBio 5: e01114–01114.

